# THiCweed: fast, sensitive detection of sequence features by clustering big data sets

**DOI:** 10.1101/104109

**Authors:** Ankit Agrawal, Snehal V. Sambare, Leelavati Narlikar, Rahul Siddharthan

## Abstract

We present THiCweed, a new approach to analyzing transcription factor binding data from high-throughput chromatin-immunoprecipitation-sequencing (ChIP-seq) experiments. THiCweed clusters bound regions based on sequence similarity using a divisive hierarchical clustering approach based on sequence similarity within sliding windows, while exploring both strands. ThiCweed is specially geared towards data containing mixtures of motifs, which present a challenge to traditional motif-finders. Our implementation is significantly faster than standard motif-finding programs, able to process 30,000 peaks in 1-2 hours, on a single CPU core of a desktop computer. On synthetic data containing mixtures of motifs it is as accurate or more accurate than all other tested programs.

THiCweed performs best with large “window” sizes (≥ 50bp), much longer than typical binding sites (7-15 base pairs). On real data it successfully recovers literature motifs, but also uncovers complex sequence characteristics in flanking DNA, variant motifs, and secondary motifs even when they occur in < 5% of the input, all of which appear biologically relevant. We also find recurring sequence patterns across diverse ChIP-seq data sets, possibly related to chromatin architecture and looping. THiCweed thus goes beyond traditional motif-finding to give new insights into genomic TF binding complexity.

## 1 Introduction

Chromatin immunoprecipitation followed by sequencing (ChIP-seq) (1) is a widely-used assay for determining transcription factor binding sites (TFBS) *in vivo*. By crosslinking the in-vivo DNA-protein complexes using formaldehyde, son-icating to break the DNA, precipitating the protein of interest using a specific antibody, reversing the crosslinks, sequencing the DNA fragments and mapping them to a reference genome, a genome-wide map of TFBS with a resolution of 100-200bp can be obtained. Newer variants like ChIP-exo (2) and ChIP-nexus (3), which promise even higher resolution, are gaining popularity. Typically these assays yield hundreds or, in large genomes, thousands to hundreds of thousands of binding sites per factor per cell type (4; 5).

TFBS are generally characterised by short conserved patterns or “motifs” in the DNA sequence, commonly represented by “position weight matrices” (PWMs) (6; 7), a probabilistic representation where each position within a binding site is described by an independent categorical distribution over the four nucleotides. A key bioinformatic task is to identify these motifs, but *ab initio* motif detection using traditional tools such as MEME (8) and Gibbs samplers such as AlignACE (9; 10) and PhyloGibbs (11; 12) is a challenge on such large datasets. Additionally, it is common for factors to interact with DNA via co-factors and not directly, which means a mixture of different motifs may be found in the ChIP-seq data.

A previous program by one of us, MuMoD (13), was targeted at the second of these problems: it simultaneously and sensitively finds multiple motifs in a given dataset. Other programs such as Chipmunk (14; 15; 16), Meme-Chip (17) and Weeder (18; 19) find successive motifs sequentially, masking previously identified sites or sequences to find the next motif.

The program we describe here, THiCweed, offers both speed and accuracy in finding multiple motifs in large datasets. It does not require prior information on the number of motifs or the lengths of the motif, since its approach is based on clustering rather than traditional motif-finding, and the clustering is based on stringent statistical criteria. On synthetic data, we show that it outperforms all current alternatives greatly on speed and is close to the best current alternative in terms of accuracy. On real genomic data, it reveals an unusual complexity in the structure of sequence motifs, in particular in internal dependencies and in flanking sequence extending far beyond the core motif.

## 2 Methods

There are two components to our approach:

- First is an efficient method of divisive hierarchical clustering. Starting with one large cluster, we split it in two clusters (or three, the third consisting of poor matches to either cluster). The scoring is described below, and is based on the likelihood ratio of a sequence belonging to one or the other cluster, done iteratively starting from an initial heuristic split. We then split each new cluster into two (or three) further clusters; and proceed until no further splits are possible. For each split we apply stringent statistical criteria to accept or reject the split. Further optimizations are described in “Algorithm”.
- During this clustering process, we include shifts and reverse complements of individual sequences to find optimal clusters. This is implemented by considering fixed-sized “windows” of length *W*, one window within each sequence. Sequences may have variable length; we permit up to half the window to lie outside the sequence, with the missing nucleotides scored as N’s, so that for each sequence of length *L*, 2*L* configurations (*L* window positions and two orientations) are considered and the optimal window chosen. The default choice of *W* is one-third the median sequence length, that is, much longer than a typical TF motif. whose positioning and orientation is sampled. This, it turns out, constitutes an effective and fast implementation of an *ab initio* motif finder on large ChIP-seq data sets, in addition to detecting the variations in motif and sequence context alluded to in the previous point. THiCweed can also be used on sequences that have been previously aligned by a “feature” (motif) to discover additional motifs/complexities, by disabling shifts and reverse complements, similar to the program NPLB (20; 21), but we don’t discuss this use here.

Our divisive clustering is in contrast to typical (agglomerative) hierarchical clustering, where individual data points are formed into clusters, requiring *O*(*N*^3^) or at best *O*(*N*^2^ log*N*) time for *N* data points. We call our approach “Top-down hierarchical clustering”; and since its purpose is to weed out “signals” in ChIP-seq peaks, we call the program “THiCweed”. (We considered “THC-weed” but it may confuse search engines.)

### 2.1 Algorithm

#### 2.1.1 Top-down hierarchical clustering

The algorithm and a typical run through it are portrayed in figure 1 and described below. We first take the simpler case of input data that has been pre-aligned with all sequences of the same length, where we don’t consider shifts and reverse-complements of sequences. The steps are as follows:

1. Initialize with one cluster containing all sequences.
2. Split every current cluster *C* (initially just one cluster), into two clusters *C*_1_ and *C*_2_, using scoring and significance criteria described below. Sequences not consistently clustering with either *C*_1_ or *C*_2_ (as described below) are concatenated into a third cluster *C_p_*. In each round, all these unclustered sequences from each division are concatenated into one cluster.
3. After every two iterations of step (2), if the current state has more than two clusters, reassign the poor-scoring sequences (sequences whose likelihoods in their current cluster are low) to the “best” available cluster.
4. Repeat from (2), until no new clusters are formed and no reassignments are made.

The user may specify a maximum number of desired clusters, and if the number of clusters at the end is greater than this, a dendrogram of current clusters is constructed and closest leaves are joined until the number of clusters is sufficiently reduced.

#### 2.1.2 Scoring

Only windowed portions of sequences are scored. Let the window length be *W*. Consider a cluster *C* with *N* sequence windows in it, *S*^1^*,S*^2^*,…,S^N^*. The probability of seeing this data if all these windows were drawn from the same PWM model is

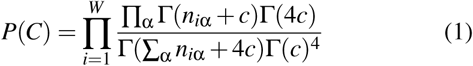

where *n_i_*_α_ is the number of times nucleotide α appears in column *i*, and *c* is a pseudocount (0.5 by default). If the cluster contains a single sequence, this expression reduces to (¼)^*W*^.

The likelihood that a sequence window *S* is sampled from the same PWM as sequences in a cluster *C* that contains *N* seqs is

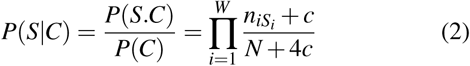

where *S_i_* indicates the *i*’th nucleotide in sequence window *S*, and *n_iSi_* is the number of occurrences of that nucleotide at position *i* in the cluster.

When splitting a cluster, an initial split is made by ranking each sequence by its likelihood of belonging to that cluster, and moving the “best” 25% to another cluster. Then sequences are selected in random order, removed from their current cluster and re-assigned to the more likely cluster, considering all possible window choices (position and orientation) within the sequence during the reassignment, until no further reassignments are made.

The significance of the split is assessed using two criteria. First, we demand that the ratio of the likelihoods of the two clusters, to the likelihood of the unsplit cluster, as calculated from equation 1, exceed a threshold, calculated from the LLR of two columns being cleanly separated in nucleotide composition. That is, suppose the two clusters consisted of random sequences, and were split on a single position – say, one cluster contained only A or C in that position, the other only G or T – while the nucleotides at all other positions are evenly distributed. This is not a significant split (it is always possible to do this, or better, for any cluster). Call the log likelihood ratio in this case *L*_1_. However, if the clusters differed in this manner in two positions – one cluster contained only A or C in those two positions, the other only G or T – this would be significant. Call the log likelihood ratio of this split *L*_2_. We demand the LLR of the split performed be equal to at least *L*_1_ +*T*(*L*_2_ *−L*_1_) where *T* is a parameter set to 0.4 by default (supplementary information). *L*_1_ and *L*_2_ can be calculated quickly using equation 1.

**Figure 1:**
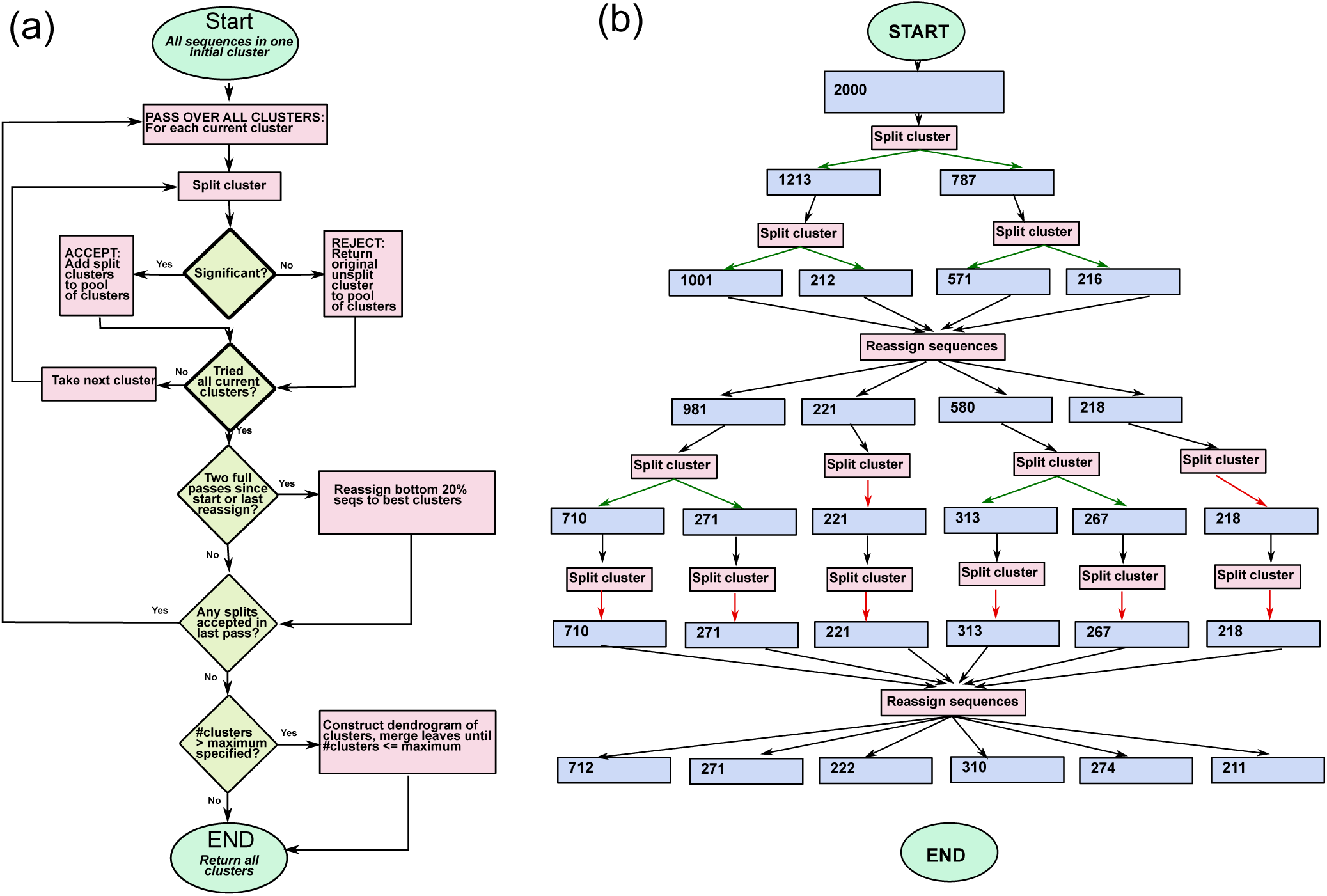
(a) Flowchart for the hierarchical clustering algorithm. The initialization is with all sequences in one cluster. At every pass, an attempt is made to split every current cluster. Splits are accepted or rejected based on significance. Every two passes, a reassignment of low-scoring sequences to the best available cluster is made. When a pass has ended with no splits being made, the program terminates returning the current clusters. (b) A possible run for an input of 2000 sequences. The blue boxes represent cluster sizes, green arrows from “Split Cluster” boxes indicate successful splits and red arrows indicate unsuccessful splits. Each horizontal row of “split cluster” boxes represents one pass.

Second, we demand that the splits be reproducible. using the following approach: we perform the split four times with four random initializations. With the resulting four pairs of clusters, we demand that at least three of the six pairwise cluster comparisons that result have an adjusted Rand index (“ARI”) (22) greater than a threshold *r* (by default 0.2). An ARI of 1.0 indicates perfect agreement while random clusterings would have ARIs close to zero. If the three pairwise comparisons between the first three splits each exceed *r*, the fourth split is not performed. If the split is accepted, the three pairs of clusters resulting from the three splits are identified based on majority membership, and sequences that failed to be consistently clustered by this criterion (that is, did not cluster in the same way according to this association) are put in a third cluster.

Splits that fail one of these two significant criteria are rejected, that is, the split clusters are joined again and returned to the pool. Both parameters *T* and *r* are user-adjustable. The default values were chosen based on benchmarks on synthetic data, as discussed in supplementary information.

When reassigning sequences (step 3 of the algorithm), we consider the poorest 20% of the sequences (measured by their likelihoods in their current clusters). For each sequence *S*, we first remove it from its current cluster, then calculate *P*(*C′*) for each available cluster *C* where *C′* = *C* + *S*, using the above formula, and add it to the best cluster. In practice, on average 4% and at most about 10% of the sequences considered in this step get reassigned.

### 2.2 Benchmarking: synthetic data

We generated synthetic datasets consisting of sequences of length 100bp each, with motifs drawn from random PWMs placed within the central 40bp of these sequences, and otherwise random (each nucleotide having probability 0.25). The PWMs had columns sampled from Dirichlet distributions with uniform hyperparameter *c* (ie, each column *v* denoting the probability distribution over the four bases A, C, G, and T, was independently sampled from the distribution 
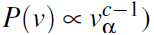
. Drawing from a Dirichlet distribution with a low value of *c* is more likely to result in a probability distribution that is highly skewed, ie is different from a uniform 0.25 probability per base. This skewness reduces with increase in *c*, a high value of *c* making the motif less distinguishable from background. Five datasets were generated with *c* = 0.1, 0.2, 0.3, 0.4, 0.5. Each dataset consisted of 20 files, with each file having sequences containing between 2 and 5 distinct motifs (one motif per sequence), the motifs drawn from PWMs of a “core” width of 5–10 bp and a tapering “flank” to a full width of 10–20bp (to reflect what is often in real data, as described below). The core positions were drawn from Dirichlet distributions with the hyperparameter *c* as described above, while the flanks tapered off rapidly from the core *c* to a hyperparameter of 20 (essentially a uniformly random vector). The performance of the programs and therefore the conclusions do not change when the flank is omitted (not shown).

Each sequence contained one motif, and each dataset contained motifs drawn from a small number of PWMs. The number and lengths of PWMs were varied across datasets for each *c*, but the distribution of numbers and lengths was the same for different *c*’s. Figure 2 shows synthetic motifs for *c* = 0.1, 0.3 and 0.5, all with a core width of 6bp and a full width of 20bp.

**Figure 2:**
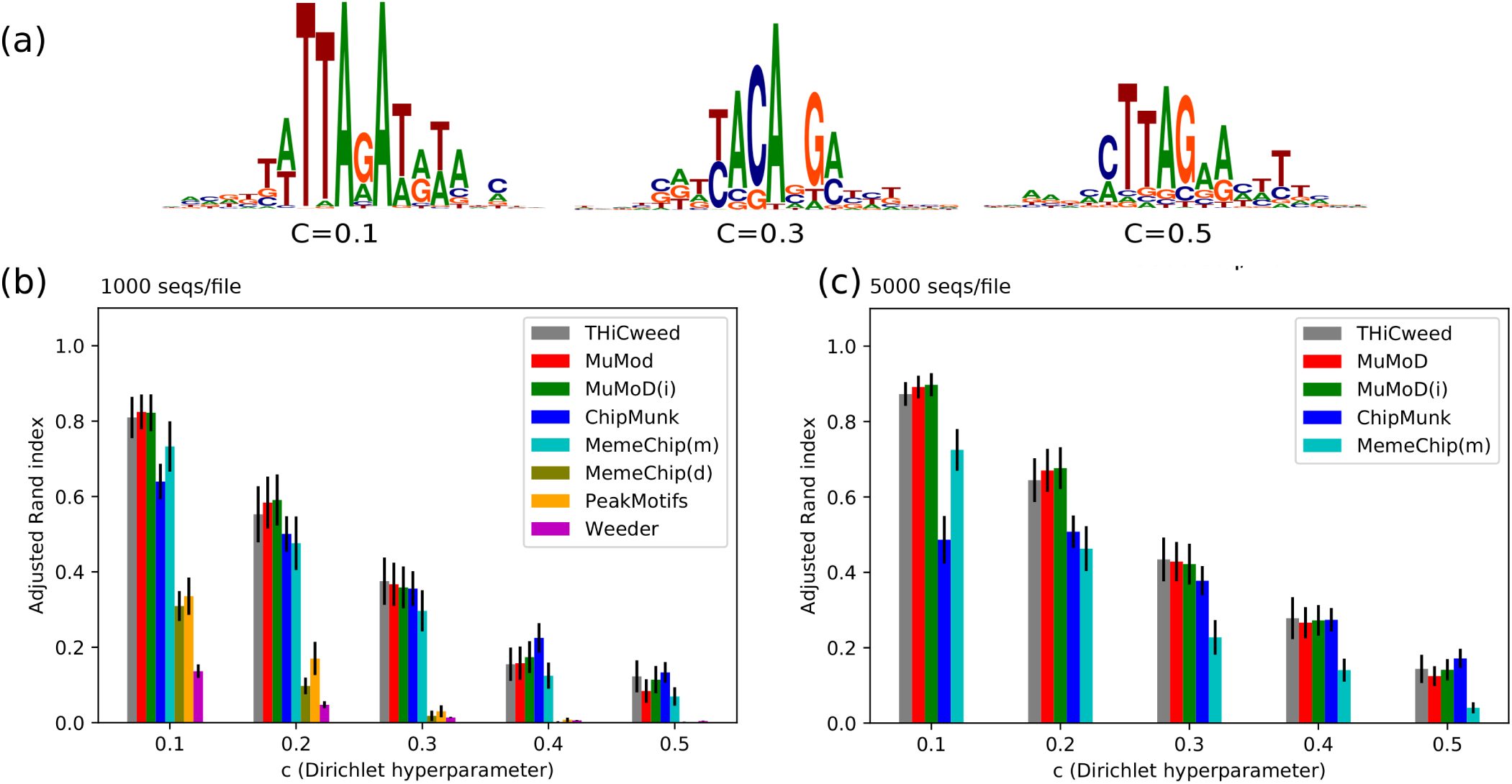
(a) Examples of embedded synthetic motifs. In this case all these have core widths of 6bp and full widths of 20bp, which are common to corresponding files in all datasets. The PWMs are sampled from different values of *c*, which varies from the indicated value in the core to a large value of 20 at the periphery. This is intended to model the appearance of motifs observed in real data. (b) and (c): Adjusted Rand index (higher is better) of predicted clustering to known clustering of synthetic data sets, containing motifs drawn from PWMs sampled columnwise from Dirichlet distributions with hyperparameter *c*. Error bars in black (standard error from 20 datasets). (b) In the case of 1000 seqs/file, THiCweed is competitive but somewhat inferior on this metric to MuMoD and ChipMunk, and somewhat superior to MemeChip (meme mode). (c) With 5000 seqs/file, comparing the better-performing programs from the previous figure, THiCweed is very close to MuMoD in performance.

THiCweed and five other programs (Peak-Motifs (23), MuMoD, Chipmunk, Meme-Chip, Weeder2) were run on these sets, in multiple-motif ZOOPS mode (zero or one occurrences of a motif per sequence). The “known” clustering of the set was the assignment of sequences to PWMs, and the “predicted” clustering for each program was the assignment of sequences to predicted motifs. The known and predicted clusters were compared using the adjusted Rand index, and the results plotted as a function of *c*. Higher ARI indicates a better match between the clusterings, with 1.0 indicating perfect agreement and 0.0 being the value expected by chance.

Two such datasets are shown here, with dataset 1 containing 1000 sequences per file, and dataset 2 containing 5000 sequences per file. The ARIs are averaged over all 20 files for each value of *c* in each dataset.

#### Commandline options

THiCweed: no additional parameters

MuMoD: Default parameters were used for the curves marked “MuMoD”. For “MuMoD(i)” the true number of motifs was specified.

ChipMunk: In all runs, the correct number of motifs was specified. The length of the motif was given as 7:20.

Weeder: Default options, but with a background frequency model derived from synthetic data.

Meme-Chip (meme): dreme was disabled with `“-dreme-m 0”`, and the known number of motifs specified with “-meme-nmotifs”, with default parameters otherwise.

Meme-Chip (dreme): meme was disabled with `“-meme-nmotifs 0”`

Peak-Motifs: Default parameters were used

#### Other notes

Despite the “filter” keyword used in the command line, Chipmunk sometimes predicts multiple motifs per sequence because it searches for matches for predicted motifs in all sequences. For computing the adjusted Rand index, each sequence was classified to the best-matching motif, as per the score reported by Chipmunk. The same was done for Peak-Motifs. In addition, sequences where no motifs were reported were assigned to an additional cluster.

### 2.3 ENCODE data

Here we used data from the ENCODE project (4; 24; 5), consisting of ChIP-seq peaks. Narrowpeak files were downloaded from the ENCODE website. 75bp flanking sequence was taken about each peak location, and repetitive regions (lowercase sequence in chromosome files downloaded from the UCSC Genome Browser (25), identified using RepeatMasker and Tandem Repeat Finder with period of 12 or less) were rejected for the purposes of this work. The cell types and ENCODE accession numbers for various factors portrayed in figures 4 through 7 are as follows, and full output is available on the web server:

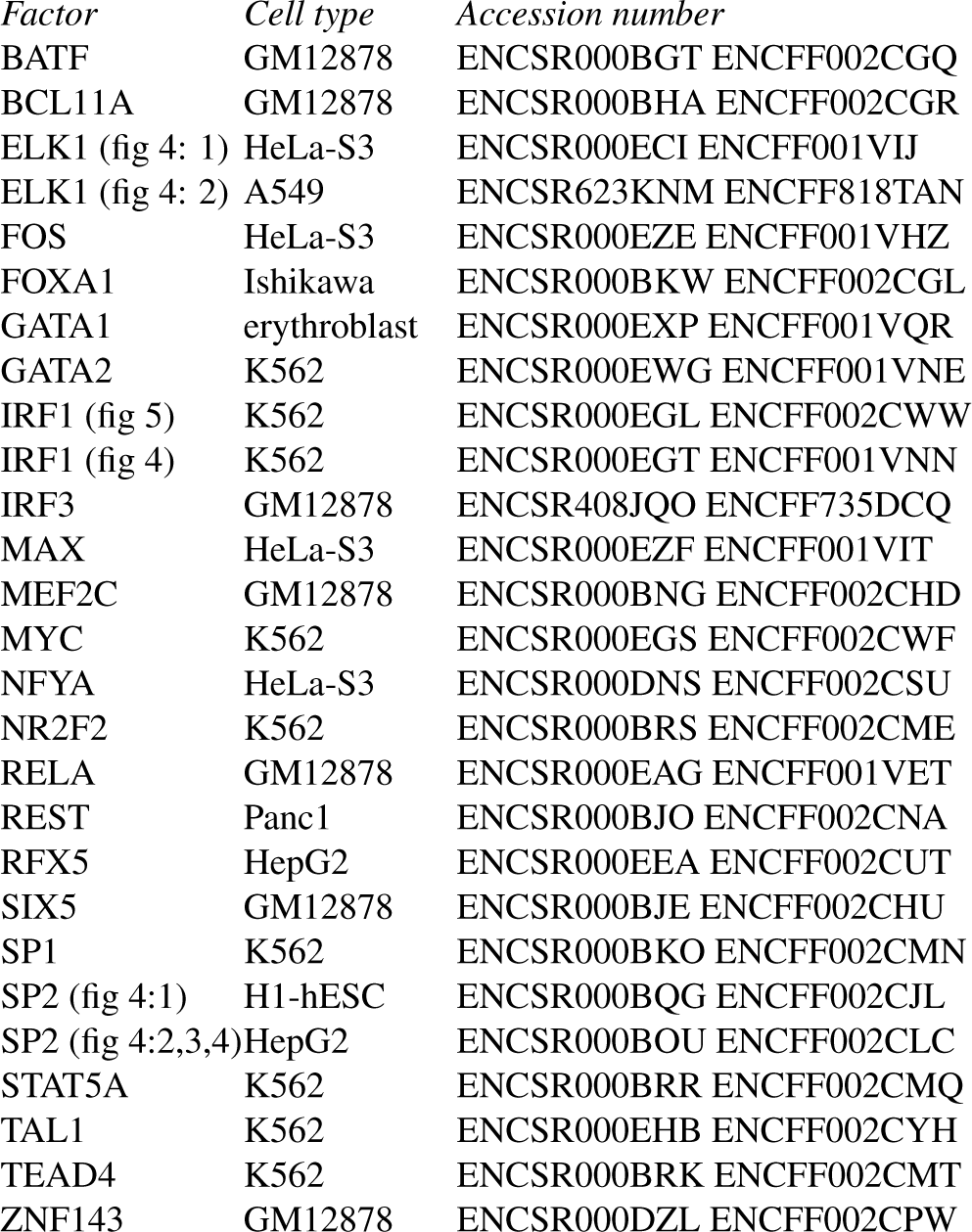

The ZNF143 clusters were compared with nucleosome positioning data in the same cell-type (GM12878) from ENCODE and PhastCons (26) phylogenetic conservation data (with other primates) from the UCSC genome site (25), distances from nearest transcriptional start sites (TSS), and DNAse-seq values from ENCODE, using custom python scripts. For TSS we used the refGene data from the hg19 release on the UCSC genome browser site.

## 3 Results

### 3.1 Synthetic data

Results for the two datasets described in Methods are plotted in figure 2 parts A,B,C for *c* = 0.1,0.2,0.3,0.4,0.5 (smaller value of *c* corresponds to sharper motifs).

In all cases THiCweed was run with default parameters, and in particular, a “window size” of 33bp or one-third the median input sequence length. As noted, it is designed to be run with large window sizes on real genomic data. Also, the stringent criteria for splitting a cluster ensure that spurious clusters are unlikely, so setting the maximum number of clusters helps only marginally (not shown). Since clusters are split according to significance criteria, there is no option to set a minimal amount of clusters.

MuMoD was run both with default parameters (“MuMoD”) and with the additional information of number of motifs (“MuMoD(i)”); the latter provides only marginal improvement. Chipmunk (in ChipHorde mode) requires the exact number of motifs to be told to it, which was done in these cases, and the range of lengths of the motif was given. Meme-Chip with its default options run the MEME motif-finder on a random subset of the input data, with inferior results. Forcing MEME for the full set improved the results, at a significant cost in running time. For comparison, we also disabled MEME entirely in favour of DREME, a heuristic approach based on regular expressions rather than PWMs. Weeder2 was run with default options but a background model derived from synthetic data, as described in Methods. With 1000 seqs/set, THiCweed is competitive with MuMoD and ChipMunk on this metric.

Only the best performers were tested with 5000 seqs/set. All programs show improved performance here, because the motif strength is maintained the same but background “noise” reduces as *N*^−½^ with increasing number of sequences *N*. But THiCweed’s improvement is sharper: it catches up with MuMoD and is largely superior to ChipMunk.

The reason for poor performance of Peak-Motifs seems to be its prediction of a very large number of motifs that are minor variations of one another. While it is hard to judge the relevance of this for real data, in the case of synthetic data these are certainly spurious, and THiCweed’s statistical criteria for splitting help it avoid this problem.

### 3.2 Running times: synthetic data

Figure 3 (a) shows running times of all the programs tested, except Peak-Motifs, for synthetic input data consisting of 200, 400, 600, 800 and 1000 sequences, each 1000bp long and containing two different motifs, each of length 10 sampled with Dirichlet parameter 0.2, in 60:40 proportion. Meme-Chip in MEME mode is an outlier: though its performance in accuracy is not very far behind other programs (figure 2, its running time would seem to disqualify it from realistic datasets (and indeed it disables MEME by default for sequence sets larger than about 600*x*100bp). It appears that, of the other programs, Chipmunk and Meme-Chip (Dreme mode) have run-times increasing roughly linearly with data size; MuMoD and Weeder running times increase superlinearly; and THiCweed’s increase is somewhat sublinear. The figure also discusses running time on genomic data from ENCODE, discussed below.

**Figure 3:**
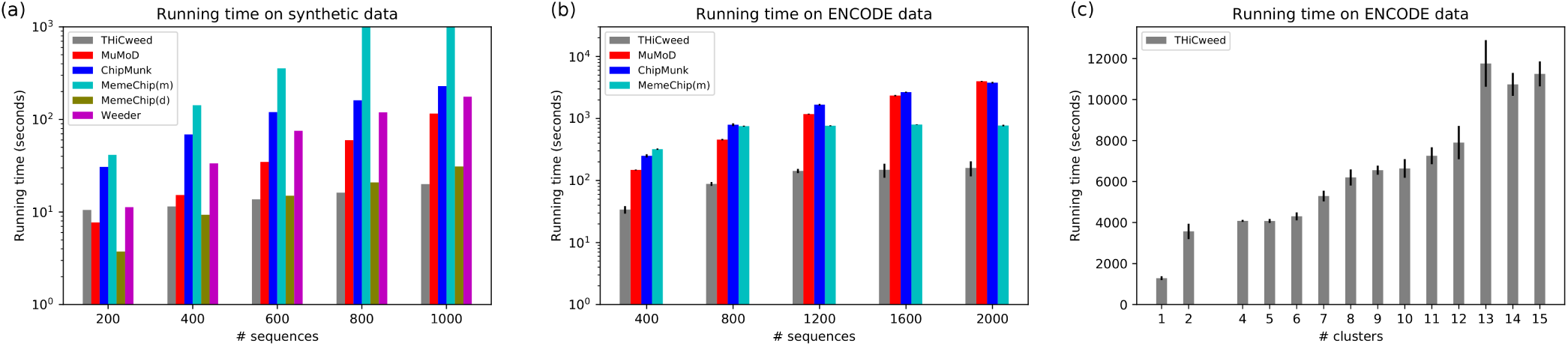
Running time of various programs. This is on synthetic data and THiCweed’s performance on real data varies significantly with the complexity of the sequence features; nevertheless, it remains on average much faster than other programs (Peak-Motifs was not tested but it is the fastest in this comparison).

### 3.3 ChIP-seq data from the ENCODE project

Running on actual genomic data yields a variety of different results depending on the factor being examined and the size of the dataset.

THiCweed has no prior knowledge of the number of different motif clusters, but by default reports a maximum of 15. In some cases far fewer are reported. Because of the statistical criteria on splitting clusters that we use, described in Methods, we believe that large numbers of clusters, if produced, are statistically significant, but THiCweed can recluster the output into smaller numbers of clusters for ease of visualization, and this is done in some cases here. Also, it works with window sizes much larger than typical motif lengths that one considers; here we used 50bp. We compare the discovered motifs to previously reported motifs from JASPAR (27; 28); the THiCweed website also includes comparisons to motifs from HocoMoco (29) and FactorBook (30).

#### 3.3.1 Ubiquitous “zinger” motifs

Hunt and Wasserman (31) observed that certain TF motifs occur repeatedly in different ChIP-seq datasets, which they termed “zingers”. In particular they identified CTCF-like, JUN-like, ETS-like and THAP11-like motifs in multiple datasets. We see all of these in our analysis of ENCODE data too (for example, the THAP11-like and CTCF motifs occur in figure 7, but several other motifs appear across multiple experiments. Figure 4 shows examples that resemble IRF1, SP2, GATA1, NFYB, REST, and a novel motif that we could not identify. Of these, SP2 and the novel motif are roughly as ubiquitous as CTCF. Both frequently co-occur with CTCF and the SP2-like motif tends to be concentrated near TSS (an example is in figure 7). We suspect a role for these in chromatin organization, a topic to be explored in future work.

**Figure 4:**
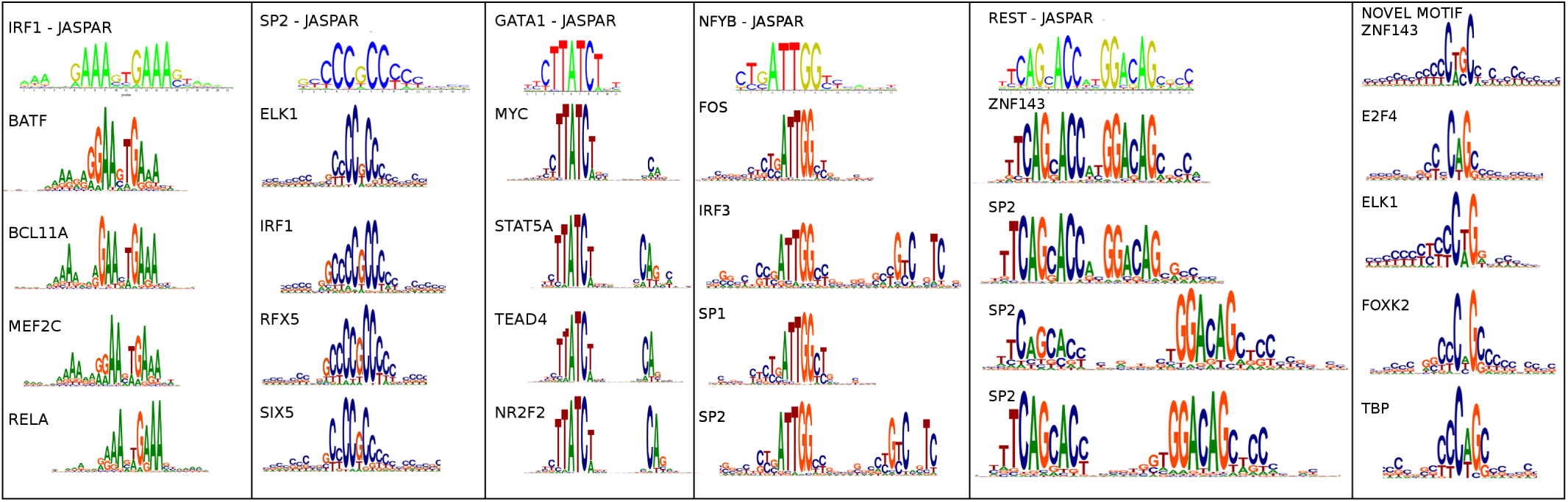
Motifs that occur across multiple chip-seq datasets, in addition to zinger motifs identified in (31). The factor for which the motif is the canonical motif according to JASPAR is indicated at the top of each column, together with the JASPAR sequence logo. Below are datasets for various other TFs where THiCweed finds the same motif.

Also noteworthy is the appearance of a secondary motif in multiple cases for the GATA-like and NFYB-like motifs; and the variable spacing of the REST-like motif. The canonical motif has two halves, TCAGCACC and GGACAG, separated by two nucleotides. But we pick up variants, previously described in (32), with longer spacing (8 and 9 bp here). Such widely spaced motifs cause problems for conventional motif-finders, but are readily picked up in our approach.

#### 3.3.2 Examples of THiCweed output

Figure 5 shows four examples of motif output. In some cases the output has been reclustered and filtered for compactness of viewing; complete results for these and many more factors are available on the THiCweed website.

We make the following observations:

- Zinger motifs are widespread here. The SP1-like motif that we documented above occurs in IRF1 and NFYA. The unidentified motif in the previous section appears in REST and FOXA1. CTCF occurs in NFYA and FOXA1. ETS-like occurs in IRF1.
- The canonical motif for IRF1 occurs in two clusters, one of which has an additional poly-T tail.
- Similarly, the canonical motif for NFYA appears in three clusters, one of which also exhibits a weak secondary motif to the left.
- The canonical REST motif occurs as a closely-spaced dimer (4th cluster), partial closely-spaced dimer (5th cluster), monomer (3rd cluster) and a widely-spaced dimer (2nd cluster). All of these variants also occur in THiCweed output for SP2 (figure 6) suggesting an interaction between SP2 and REST. The widely-spaced dimer is not picked up by other motif finders.

**Figure 5:**
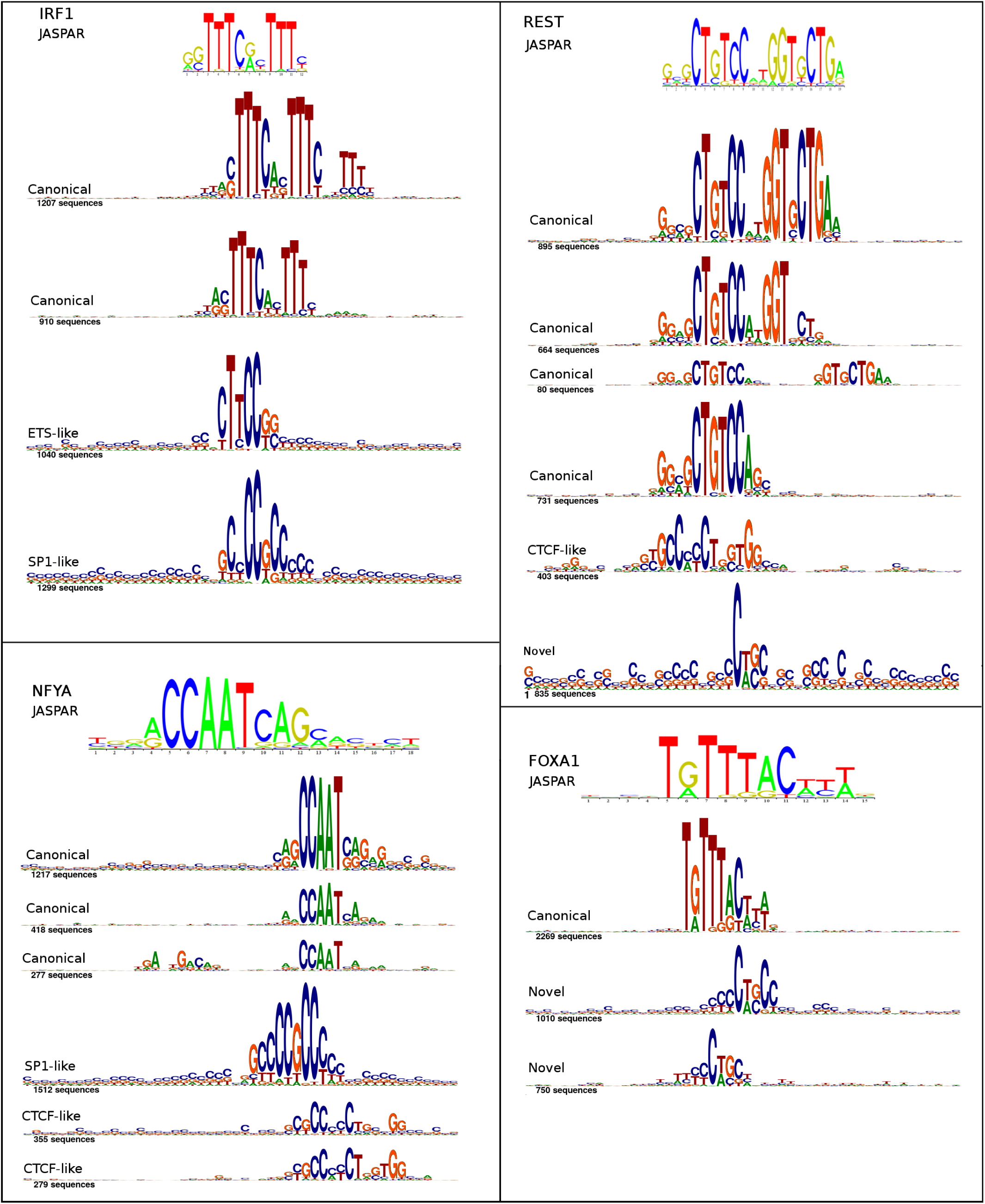
Sample THiCweed output on four ChIP-seq datasets: IRF1 (5543 peaks), NFYA (4497 peaks), REST (3998 peaks), FOXA1 (4029 peaks). Not all output clusters are shown here. The full output is available on the THiCweed website.

**Figure 6:**
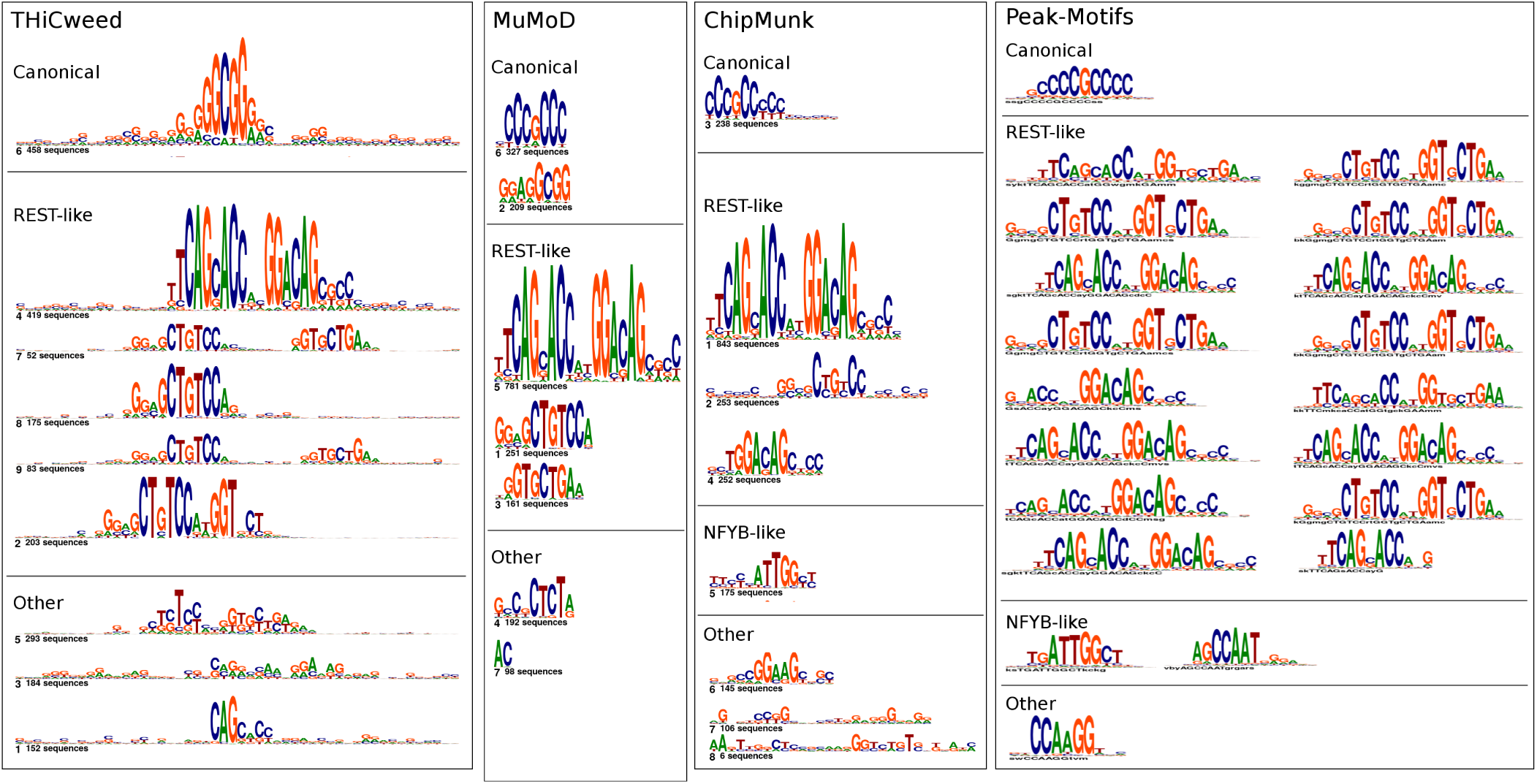
Comparison of clustering of 2,019 peaks for SP2 by THiCweed, with motifs found by three other programs

A much larger collection of THiCweed output on ENCODE factors is available on the website. Features similar to those noted above are ubiquitous.

#### 3.3.3 Comparison with other programs

Figure 6 compares the output of THiCweed with three other programs. All programs pick up the main motif (though with varying numbers of instances). All also pick up the REST motif, but only THiCweed picks up the widely-spaced version in one piece. THiCweed also seems to reveal a larger surrounding-sequence context in many cases, notably for the SP1-like motif which generally occurs in a CG-rich background. Peak-Motifs identifies a very large number of motifs, most of which appear to be minor variations of the main motif. This may explain the poor performance of Peak-Motifs on our synthetic benchmark: the adjusted Rand index would penalize breaking up clusters into smaller clusters.

#### 3.3.4 Biological relevance of these clusters

We typically find several different motifs, variants of a motif, and a few apparently uninformative clusters in THiCweed runs. Biological significance to these are suggested on comparing other genomic features such as phylogenetic conservation (via PhastCons scores (26) from the UCSC Genome Browser (25)) and nucleosome occupancy and DNAse-seq data (from ENCODE (4)). Figure 7 compares each of 8 clusters for ZNF143 with a plot of conservation, nucleosome occupancy (in an extended region of 1000bp on each side), distance to the nearest TSS, and DNAse-seq values. Cluster 6 (SP2-like motif) tends to be concentrated close to TSSs (mostly within 1000bp – a pattern we see consistently), shows little phylogenetic conservation, and no sign of nucleosome positioning. Cluster 1 (a motif resembling THAP11, identified in (31) as a zinger motif), too, is concentrated near TSSs; it too shows little effect in nucleosome positioning, but is strongly conserved. Cluster 7, resembling the REST motif, is spread away from TSSs, is phylogenetically conserved, and has an effect on nucleosome positioning (which we observe in other datasets where this motif occurs).

Cluster 8 seems uninformative, but it appears concentrated near the TSSs (within about 5000bp), which would likely not happen if it consisted only of random unclusterable sequences left over from the other clusters.

The remaining clusters are variants of the CTCF motif; cluster 5 includes the previously documented “M2” motif. Cluster 3 appears different from other CTCF clusters in that it occurs in a GC-rich background, is more concentrated near TSS (mostly within about 10000bp), appears a little less conserved and a little less effective at nucleosome positioning, with more open chromatin as shown by DNAse.

### 3.4 Running times: ENCODE data

Figure 3 (b) shows the results of THiCweed, MuMoD, ChipMunk and Meme-Chip (MEME mode) on real ENCODE data, consisting of 400–2000 random samples from a set of CTCF ChIP-seq peaks (dataset ENCFF001USS). The results are similar to on the synthetic data, except that, somewhat surprisingly, Meme-Chip is faster than MuMoD and ChipMunk on larger datasets.

Figure 3 (c) shows the running time of THiCweed as a function of the number of clusters found, on 92 ChIP-seq datasets each consisting of 27000–33000 peaks, across multiple TFs and cell lines. The running time increases with the number of clusters, but somewhat sublinearly. On such realistic ChIP-seq datasets, THiCweed’s running time is about two orders of magnitude less than MuMoD, which can take days, and is also much faster than all other programs tested. Meme-Chip uses the MEME step on only a small fraction of the input sequences; and Weeder2 learns motifs from a small fraction of the sequences and uses those to analyse the rest (19). THiCweed processes the majority of files of this size in under two hours, with interesting and biologically relevant results.

## 4 Discussion

Motif-finding in large datasets produced by ChIP-seq and similar experiments is a qualitatively different problem in complexity from what traditional motif-finders are used to handle. Additionally, one could liken the problem of finding rarely-occurring motifs to finding a needle in a haystack. We view THiCweed’s approach as “sequence feature analysis” (over large windows) rather than “motif-finding” (detection of short patterns). Our novel clustering algorithm can comfortably handle tens of thousands of sequences at a time, and with significant heterogeneity in motif content. It successfully picks up biologically relevant motifs even when they occur in fewer than 5% of the input sequences, such as the REST-like motif in ZNF143 (cluster 7 in figure 7). Its large window size enables it to also pick up secondary motifs like the M2 CTCF motif in the ZNF143 data (figure 7), the widely-spaced dimer in SP2 and REST (figures 5 and 6), and peripheral features such as an overall CG-richness in some motifs (eg CTCF-like cluster 3 in figure 7). The significance criterion used for splitting, and the differences in biological parameters in figure 7, suggest that these differences are important and are not artifacts.

**Figure 7:**
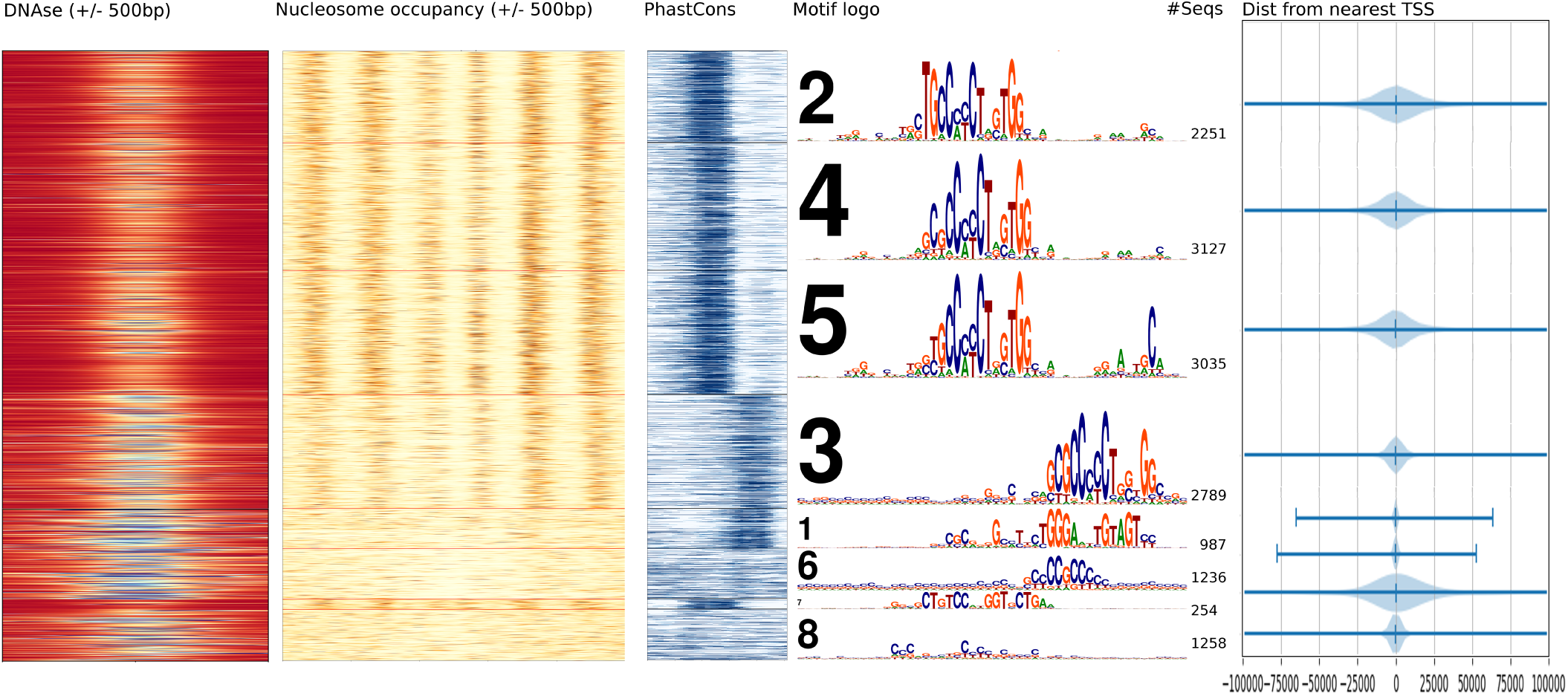
Comparison of sequence clusters of 14,937 ZNF143 ChIP-seq peaks with DNAse-seq values (colour scale: blue=open, red=closed), nucleosome occupancy (colour scale: white = 0, brown = 5+), PhastCons conservation score (colour scale:white=0, dark blue=1), and distance from nearest TSS, suggesting connections between the motif structure in different sequence clusters, biological function, evolutionary conservation pressure, nucleosome positioning and open/closed chromatin.

Uniquely among the programs we have tested, THiCweed achieves its combination of speed and accuracy without resorting to heuristics in scoring (as DREME and Chipmunk do, using regular expressions and “seeding” respectively) and without resorting to training on a small subset of the sequences (as Weeder does). THiCweed’s clustering algorithm is stochastic, but is essentially similar to an iterated *K*-means clustering with *K* = 2, with significance criteria to avoid spurious splits. Instead of invoking pairwise distances and calculating a centroid, however, we calculate multinomial likelihoods correctly within the limitations of the PWM assumption. The clustering algorithm and wide-window approach ensures that little or no prior information is required to run the program: significant short motifs can be found inside longer windows by eyeballing, but other relevant sequence features can be picked up too.

A possible shortcoming is that within THiCweed’s framework, only one motif occurrence per sequence will be detected (unless two motifs co-occur with a restricted spacing, as in the extended REST motif and the secondary M2 CTCF motif). Sequences that match no dominant motif may end up in a relatively uninformative cluster such as cluster 8 in figure 7. One may ask whether, in clusters that do not match the canonical motif, the motif nevertheless occurs elsewhere in some of the peaks in additional to the non-canonical motif in the cluster. We checked for this possibility exhaustively using FIMO (33), with a *q*-value threshold of 10^*−*3^, and found that, out of 93 TFs that have a PWM in JASPAR, only 25 reported any sequences at all that were not clustered with canonical motif matched but that nevertheless showed hits for the JASPAR motif. These too showed matches in very few cases; the exceptions were SP2 and EGR1, both of which have GC rich canonical motifs, which reported matches in about 15% and 14%, respectively, of such sequences. It would therefore seem that the “missing” of canonical motifs because of occurrence of other strong motifs within ChIP-seq peaks is not a common concern in practice.

At the moment, THiCweed runs on a single CPU core, but significant speedups are possible by parallelization.

In cases where there is a profusion of similar but slightly different motif patterns as well as an occurrence of many different motifs (as in the ZNF143/CTCF case), it appears that the differences may have biological significance, as reflected by nucleosome positioning and phylogenetic conservation. We plan to explore this, and the significance of some of the novel zinger motifs, further in a future work.

## Acknowledgements

We thank Vishaka Datta and Arvind Shankar for useful discussions. The plots in figure 7 were generated with a script written by Arvind Shankar for a different project which will be reported elsewhere.

## Author contributions

AA, LN and RS conceived the basic algorithm. AA implemented an early prototype. RS implemented the current version in the Julia language. AA, LN and RS contributed to various refinements in development. SVS and RS implemented the webserver. All authors contributed to the benchmarks described here and to the analysis of ENCODE chip-seq data. All authors read and approved the final manuscript.

**Availability:** The software is open source and available for download at http://www.imsc.res.in/~rsidd/thicweed/ under the two-clause BSD license. An online web-server is also available, linked on the above page, and can be used for modest-sized jobs.

